# Cytokine profile in human olfactory cleft mucus and associated changes in olfactory function

**DOI:** 10.1101/332395

**Authors:** Hong Wang, Cristina Jaen, Keiichi Yoshikawa, Mai Haneoka, Naoko Saito, Junji Nakamura, Nithin D. Adappa, Noam A. Cohen, Pamela Dalton

## Abstract

Multiple factors, including physical changes of the nasal mucosa and epithelium and exposure to air-borne environmental agents, appear to contribute to age-related olfactory loss. However, the molecular aspects of aging-associated olfactory loss in humans are not well understood. Although inflammation can be a significant underlying cause for olfactory impairment, whether aging increases the levels of inflammatory cytokines in the human olfactory mucosa and whether any inflammatory markers are associated with age-related olfactory loss remain unclear. Using a noninvasive method for collecting human olfactory mucus, we characterized and compared inflammatory cytokines, chemokines, and some growth factors, in the mucus collected from the olfactory cleft or the anterior nasal cavity from 12 healthy, young (18-40 years old) and 12 elderly (60-85 years old) individuals. We also hoped to identify candidate molecular biomarkers associated with age-associated olfactory loss in humans. Olfactory thresholds were obtained for two odorants and individual mucus samples were analyzed using multiplex assays for the levels of 30 cytokines. Results indicated elevated levels of certain inflammatory cytokines (IL-12, MCP-1) in olfactory mucus of the elderly, and high levels of some inflammatory factors (MCP-1, IL-8, IL-13 and VEGF) were associated with reduced olfactory sensitivity, suggesting that inflammation may play a role in olfactory decline associated with aging.

## Introduction

Age-related decreases in olfactory acuity have been well-documented, beginning around the 5th decade of life and worsening with age (1-3). Such decreases are manifested in changes in threshold sensitivity as well as the ability to discriminate and identify odors. Olfactory loss is also one of the earliest symptoms of several age-related neurodegenerative diseases, including Parkinson’s disease and Alzheimer’s disease (4, 5). Multiple factors appear to contribute to age-related olfactory loss. Some relate to changes in the nose such as atrophy of the nasal epithelium, decreases in mucosal blood flow, and impairment of mucociliary function. Replacement of olfactory epithelium with respiratory epithelium has also been noted among the elderly (6). Decades of exposure to air-borne environmental agents may also contribute to age-related olfactory loss (7). However, the molecular aspects of aging-associated olfactory loss in humans are not well understood.

Inflammation may play significant roles in chemosensory disorders (7-10). Some diseases with robust inflammatory responses in the nose, such as upper respiratory infection and chronic rhinosinusitis, are strongly associated with olfactory loss (3, 11-13). In addition, shown in animal studies, inflammation and some inflammatory cytokines induced in the olfactory mucosa inhibit neurogenesis and/or stimulate apoptosis of olfactory neurons (8, 14, 15). In the aging olfactory mucosa, cellular injuries and stress, resulting from life-long exposure to harmful chemicals, allergens, and pathogens, could stimulate inflammation (7). However, whether aging increases the levels of inflammatory cytokines in the human olfactory mucosa and whether any inflammatory markers are associated with age-related olfactory loss remain unclear.

Aging is associated with low-grade chronic systemic inflammation, characterized by elevated levels of circulating inflammatory markers (16, 17). Yet, a recent report showed that the baseline blood levels of three inflammatory markers, C-reactive protein, interleukin (IL)-6, and tumor necrosis factor (TNF), are not associated with long-term (10 years) development of olfactory impairment in older adults (53-97 years old) (18). This study suggests that inflammatory proteins in the blood may not be good indicators for the inflammatory status in the olfactory mucosa and do not show strong correlation with olfactory function. To better understand the relationship between inflammation and aging-related olfactory decline, direct measurements of inflammatory cytokines and biomarkers in the olfactory tissue could be more informative. Yet, it is challenging to obtain human olfactory tissue biopsies, because the procedure is fairly invasive. Thus, establishing a noninvasive method to assess local inflammation in the olfactory mucosa is an important step towards understanding of the role of inflammation in olfactory loss during aging.

The olfactory epithelium is covered by a layer of mucus with a depth of about 5-39 um, which serves to protect the olfactory neurons and provide a constant aqueous environment around the receptors (19). Unlike the mucus covering adjacent nasal respiratory epithelium, which is produced primarily by goblet cells and anterior nasal seromucous glands (20), the mucus in mammalian olfactory cleft is mainly from Bowman’s glands and the supporting, sustentacular cells in the olfactory tissue (21). The olfactory mucus is an important component of the olfactory system. It provides a critical interface for odorants to access olfactory receptor neurons. Odorant binding proteins in the mucus are necessary to transport lipid molecules through the mucus to the receptors (22-24). Olfactory mucus also plays other important roles in olfaction, such as buffering and removal of harmful environmental toxins and odorants, participating in olfactory epithelium turnover, and orchestrating local immune responses to pathogens. Consistent with these functions, various enzymes, growth factors, and cytokines are found in the olfactory mucus (25-27). It is possible that the molecular components of the olfactory mucus could provide biomarkers for the molecular, cellular, sensory processes occurring in the olfactory system.

The goals of this study were to (1) evaluate a noninvasive method for collecting human olfactory mucus, (2) characterize and compare the profiles of inflammatory cytokines, chemokines, and some growth factors in the mucus collected from the olfactory cleft (OC) or the anterior nasal cavity (ANC) from healthy young adult and elderly subjects, and (3) identify candidate molecular biomarkers associated with human olfactory loss during aging. We collected olfactory mucus from healthy, young (18-40 years old) and elderly (60-85 years old) individuals, using a noninvasive collection technique. In order to determine the degree to which mucus collected from the olfactory cleft truly differs from mucus collected more anteriorly, we collected both types of mucus from each individual. Mucus was analyzed using the sensitive Luminex multiplex platform (28) for the levels of 30 cytokines and growth factors. The global protein contents of the mucus were also analyzed by a shot-gun proteomics ANCproach (see companion article, Yoshikawa et al.). Our results show that the majority of the analyzed cytokines and growth factors can be detected in the collected OC and ANC mucus from at least 50% of the subjects. The levels of interleukin-12 (IL-12, p40/p70) and monocyte chemoattractant protein-1 (MCP-1, or CCL2) are significantly elevated in the olfactory mucus of elderly subjects. In addition, the levels of MCP-1, IL-8 (CXCL8), IL-13, and vascular endothelial growth factor (VEGF) in the olfactory mucus are significantly, inversely associated with the combine threshold sensitivity to phenyl ethyl alcohol (PEA) and n-butanol. These results indicate that the levels of certain inflammatory cytokines are elevated in the olfactory mucus of the elderly, and high levels of some inflammatory factors are associated with reduced olfactory sensitivity. Our study suggests that inflammation may play an important role in olfactory decline associated with aging.

## Results and Discussion

### Cytokine profile in the nasal mucus from healthy adults: high levels of IL-1RA and IL-8

Nasal mucus from the OC and ANC was collected using a noninvasive method. Thirty cytokines and growth factors (Supplemental Table 1) were analyzed by multiplex bead assays using the Luminex platform. The majority of the 30 cytokines and growth factors can be detected in at least half of the mucus samples from either the OC or ANC (Supplemental Table 1 and 2). However, the levels of the 30 cytokines and growth factors in the nasal mucus were dramatically different.

IL-1 receptor antagonist (IL-1RA) showed the highest levels among the 30 analytes in both the ANC and OC mucus (Fig. 1A and 2A). The means of IL-1RA levels in the ANC and OC mucus from the young subjects were 833176.5 pg/ml and 1409488.6 pg/ml, respectively (Supplemental Table 1). IL-1RA was the only cytokine among the 30 analytes in the OC and ANC mucus that was detected by the shotgun proteomics approach, which preferentially detected proteins at high concentrations and/or with high molecular weight (see companion article, Yoshikawa et al.). High levels of IL-1RA in nasal secretions or lavages have been reported previously and are consistent with our results (29, 30). The cytokine that showed the second highest levels in the nasal mucus was IL-8 (Fig. 1A and 2A). The means of IL-8 levels in the ANC and OC mucus from the young subjects were 81742.5 pg/ml and 44309.1 pg/ml, respectively (Supplemental Table 1). Relatively high levels of IL-8 in nasal lavage or mucus were reported previously by us and others (27, 31). Conversely, several cytokines, including IL-1β, IL-2, IL-4, IL-5, IL-15, IL-17, and TNF, were not detected in a significant percentage of mucus samples, and if they were detected, their levels were in the low pg/ml range.

**Figure 1.**
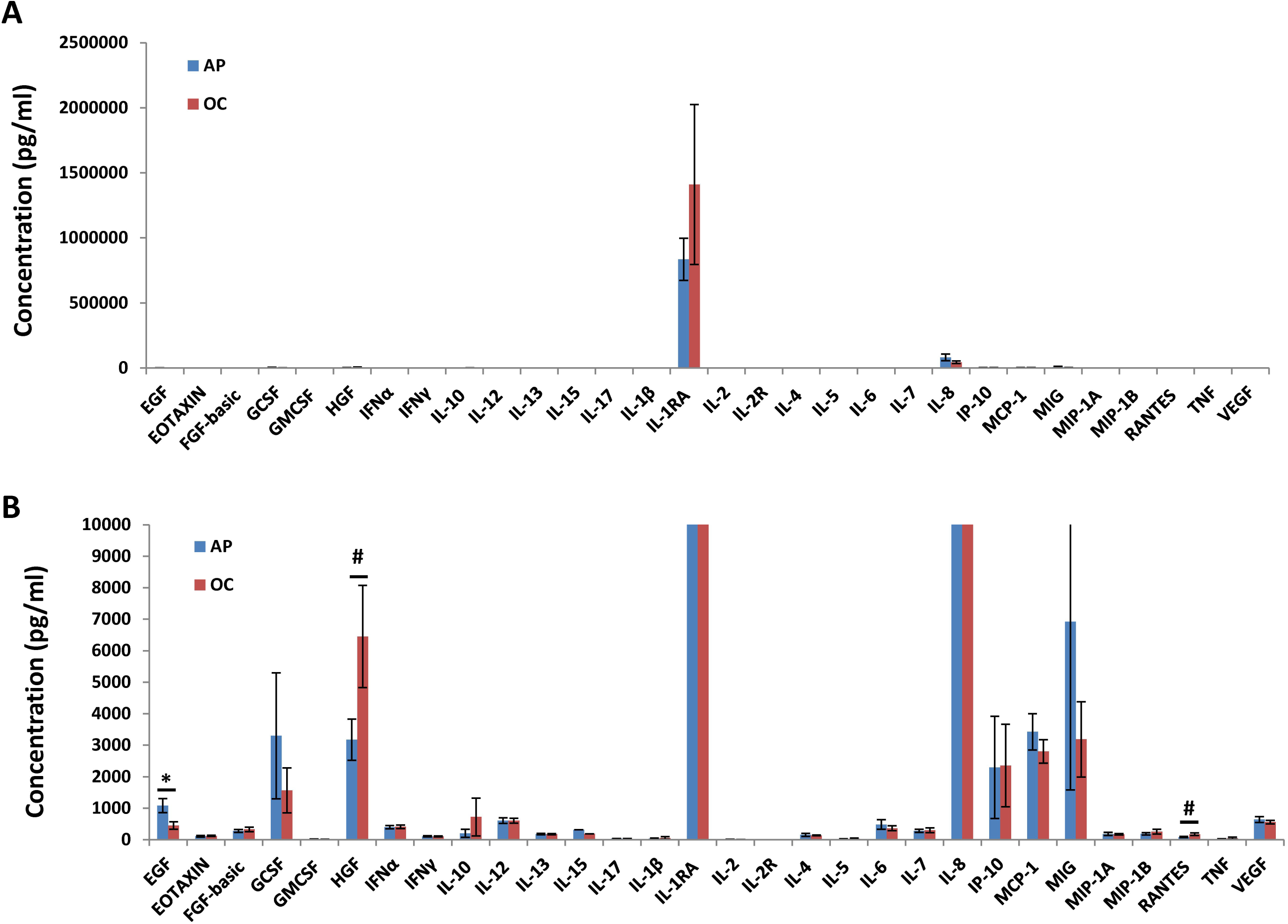
Regional differences in the levels of cytokines and growth factors in the nasal mucus from young subjects. Data were from Luminex multiplex cytokine assays, showing the levels of 30 cytokines and growth factors in the ANC and OC mucus. (A) Cytokine and growth factor levels showing under large concentration scales to highlight the levels of the two most abundant cytokines, IL-1RA and IL-8. (B) Cytokine and growth factor levels showing in the low concentration range. Data are mean ± SEM. N = 12. * *p* < 0.05. # *p* < 0.08.

**Figure 2.**
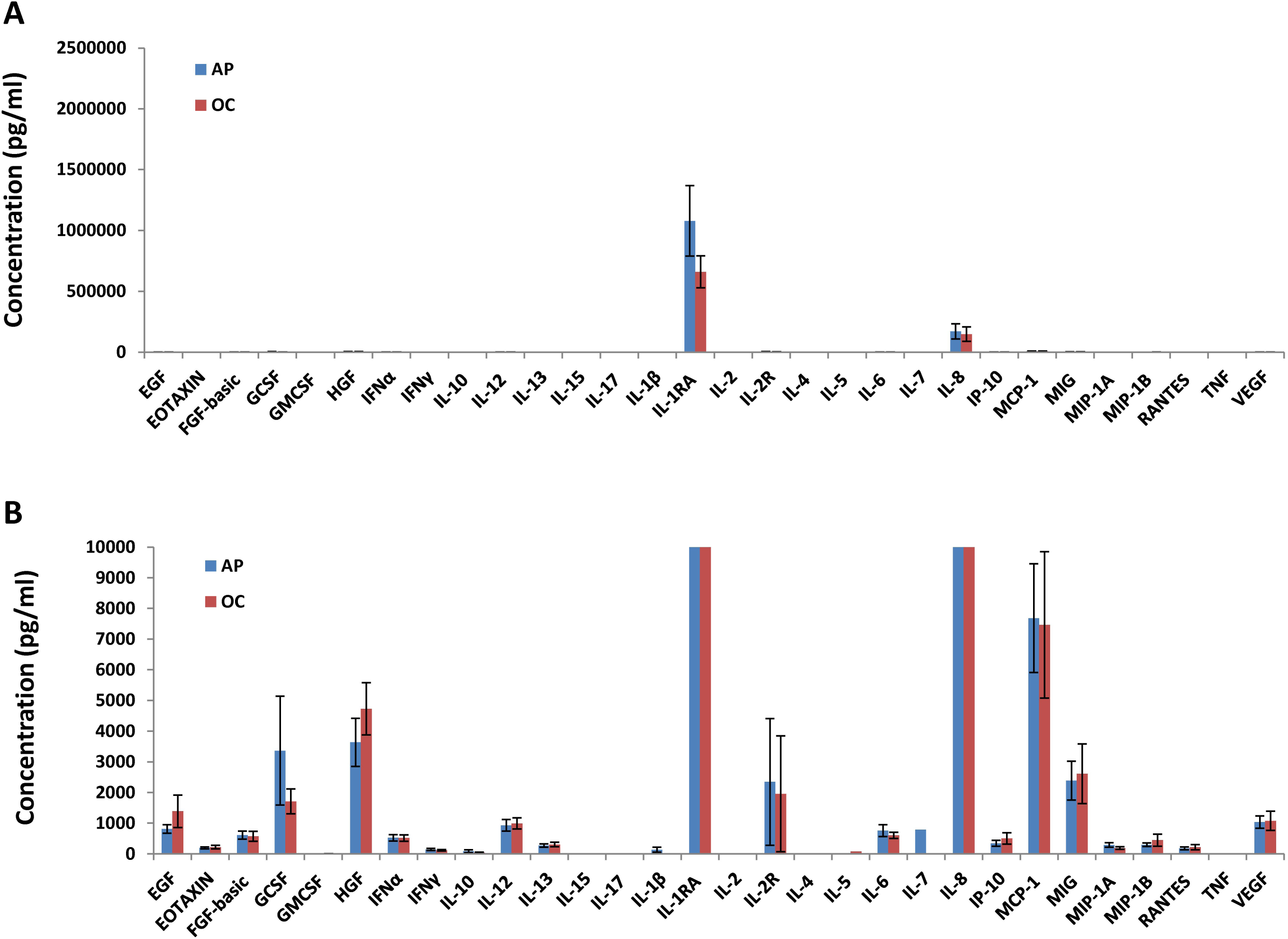
Cytokine profile in the nasal mucus from elderly subjects. Data were from Luminex multiplex cytokine assays, showing the levels of 30 cytokines and growth factors in the ANC and OC mucus. (A) Cytokine and growth factor levels showing under large concentration scales to highlight the levels of the two most abundant cytokines, IL-1RA and IL-8. (B) Cytokine and growth factor levels showing in the low concentration range. Data are mean ± SEM. N = 10-12.

Considering that the subjects were healthy, without any subjective or objective evidence of sinonasal inflammation, the low levels of some of these cytokines in the mucus are not surprising. On the contrary, the high levels of IL-1RA and IL-8 suggest that these two cytokines may play special roles in the nasal cavity and the olfactory mucosa. IL-1RA is a naturally occurring, endogenous IL-1 receptor antagonist and plays important anti-inflammatory roles in multiple tissues. Deficiency of IL-1RA, due to loss-of-function mutations, results in systemic auto-inflammatory diseases (32). IL-1RA polymorphism is also associated with chronic rhinosinusitis and asthma (33, 34). The IL-1 receptor IL-1R is ubiquitously expressed in virtually all cell types and is a key signaling receptor for the IL-1 family of inflammatory cytokines (35). As an endogenous IL-1 receptor antagonist, IL-1RA may play critical roles in shutting down IL-1R signaling in the nasal respiratory tissues and the olfactory mucosa to inhibit allergen- and/or infection-induced inflammatory responses (7). The high levels of IL-1RA are likely needed to protect the tissues from inflammation-related injury. IL-8 is a major inflammatory mediator and a key chemotactic factor for recruiting and activating immune cells, especially neutrophils, which play important roles in innate immunity (36). High levels of IL-8 in the nasal mucus from healthy adults suggest that immune cells are continuously directed to the nasal mucosa for immune surveillance and/or clearance even under normal conditions.

### Comparison of cytokine profiles in the nasal mucus from the ANC and OC regions: differential levels of EGF in young adults

Although the olfactory and respiratory mucus in the nose are produced by different glands and epithelial cells, a continuous layer of mucus bathes the nasal epithelium from the OC to ANC and some mixing occurs. To assess whether differences exist in the levels of cytokines and growth factors in the respiratory and olfactory nasal mucosa, the ANC and OC mucus was collected and analyzed from each subject. As shown in Fig. 1B, in young subjects, the level of epithelial growth factor (EGF) in the ANC mucus was significantly higher than that in the OC mucus (*p* = 0.028, Supplemental Table 1). In addition, the *p* values for two other cytokines, hepatocyte growth factor (HGF) and regulated upon activation normal T cell expressed and secreted (RANTES), were close to be significant (*p* = 0.077 and 0.059, respectively); and in these cases, the levels of HGF and RANTES were higher in the OC mucus than in the ANC mucus (Fig. 1B and Supplemental Table 1). The differential levels of EGF, HGF, and RANTES in the ANC versus OC mucus are highly consistent with a previously published study on transcriptome analyses of human nasal respiratory and olfactory epithelia (37). The relative levels of the mRNA transcripts for EGF, HGF, and RANTES in the respiratory and olfactory tissues are in the same trends as those in our data on protein levels of these factors in the ANC and OC mucus. Interestingly, the differences in these cytokine levels between the ANC and OC mucus from the young subjects were much reduced in the mucus from the elderly subjects (Fig. 2B and Supplemental Table 2). In fact, the mean of EGF levels in the ANC mucus was lower than that in the OC mucus from the elderly subjects, which was the opposite as shown in the young subjects.

These results indicate that, even though some mixing of the ANC and OC mucus occurs and the levels of the majority of the cytokines and growth factors were comparable in the ANC and OC mucus, the levels of some other factors can be significantly different in the ANC versus OC mucus. The differential levels of these factors could be the result of differential mRNA expression in the respiratory versus olfactory tissues in the nose (37). These results also argue that the OC mucus is a better choice of sample source than the ANC mucus for studying the molecular and cellular processes in the human olfactory epithelium.

Both EGF and HGF are important growth factors that regulate epithelial cell growth, differentiation, and morphogenesis. EGF was shown to have mitogenic effects (i.e. stimulating cell proliferation) on *in vitro* cultured olfactory neuron progenitor cells and to stimulate olfactory neuron reinnervation of the olfactory bulb after tissue lesion in rats (38, 39). However, considering the level of EGF in the young OC mucus is lower than that in the ANC mucus, it is possible that too high levels of EGF may promote differentiation of respiratory rather than olfactory epithelium in humans. Interestingly, the level of EGF in the OC mucus from the elderly is much higher than that in the young OC mucus. It is reasonable to speculate that this high level of EGF may contribute to the reported replacement of olfactory epithelium with respiratory epithelium in the elderly (6). The higher levels of EGF in the elderly OC mucus could also be due to increased activities of tissue repair. HGF and its receptor were shown to be highly expressed in the developing olfactory system (40). In *in vitro* cultures, HGF can stimulate the proliferation of olfactory ensheathing cells (41). The higher levels of HGF in the young OC mucus may suggest that this growth factor could play a role in cell proliferation and survival in the olfactory epithelium. Further studies are needed to better understand the roles of EGF and HGF in human olfactory neuron generation, turnover, and survival.

### Aging-related changes in cytokine profiles in the ANC and OC mucus: increased levels of inflammatory cytokines in the elderly

To investigate whether aging changes cytokine levels in the nasal mucus, especially in the OC mucus, we compared cytokine profiles from the young and the elderly. As shown in Fig. 3, the levels of IL-12 (p40/p70) and MCP-1 were significantly higher in the elderly OC mucus than in the young OC mucus (*p* = 0.0497 and 0.047, respectively, also see Supplemental Table 3). The levels of IL-6 and IL-8 were also higher in the elderly OC mucus than in the young OC mucus (although did not reach statistically significant level, *p* = 0.068 and 0.074, respectively). All four cytokines are strong mediators of inflammation. Their increased levels in the elderly OC mucus indicate that aging is associated with higher inflammatory responses in the olfactory mucosa. Consistently, the levels of IL-10 and IL-1RA, two major anti-inflammatory cytokines, were decreased in the elderly OC mucus (although did not reach statistically significant level), suggesting a reduction of anti-inflammatory activities. Our previous studies showed that in the taste sensory tissue IL-10 deficiency leads to structural defects, probably by increasing inflammatory responses that slow down cell renewal and accelerate cell death (9, 42, 43).

**Figure 3.**
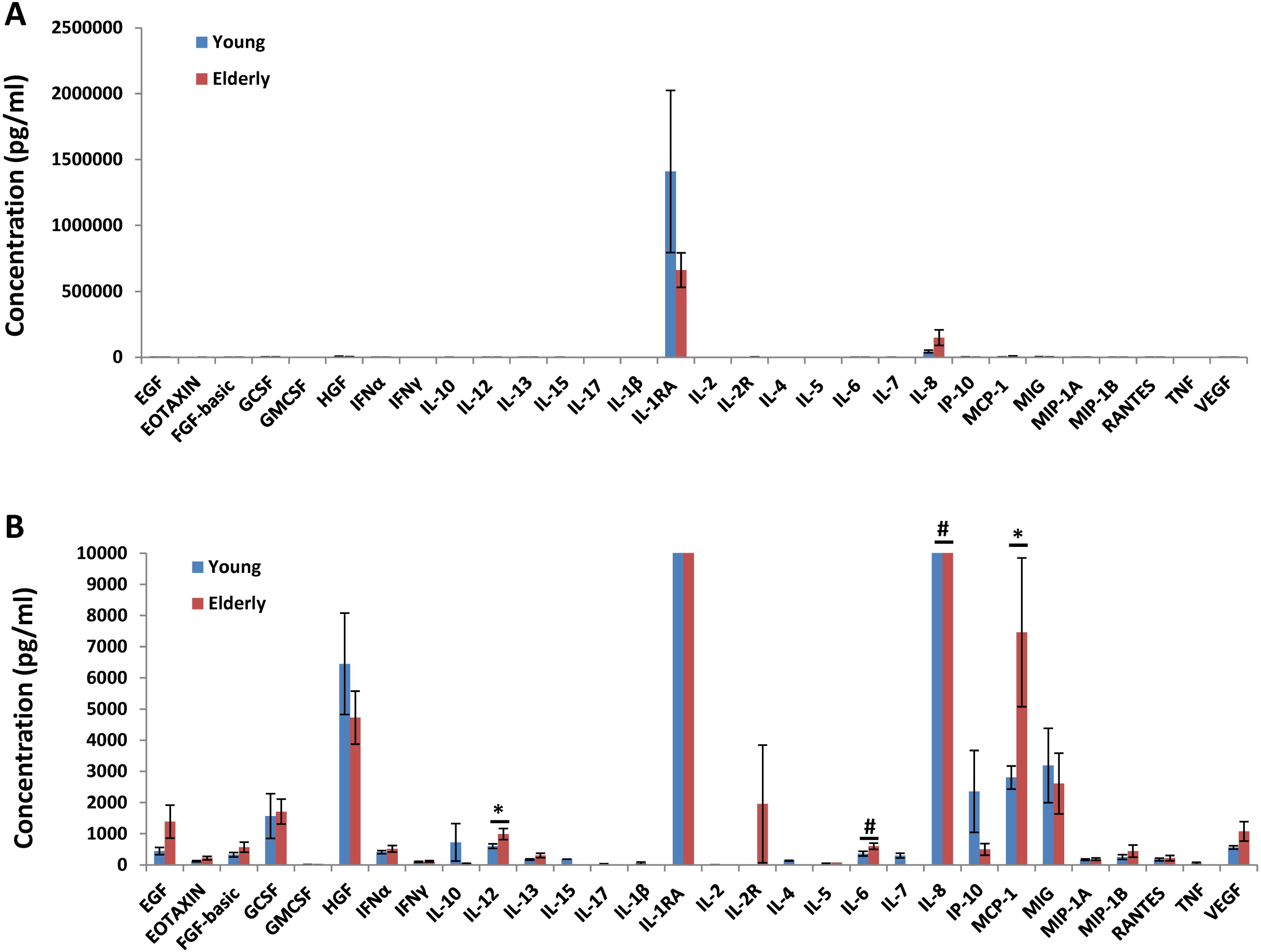
Comparison of cytokine profiles in the OC mucus from young and elderly subjects. Data were from Luminex multiplex cytokine assays, showing the levels of 30 cytokines and growth factors. (A) Cytokine and growth factor levels showing under large concentration scales to highlight the levels of the two most abundant cytokines, IL-1RA and IL-8. (B) Cytokine and growth factor levels showing in the low concentration range. Data are mean ± SEM. N = 10-12 per group. * *p* < 0.05. # *p* < 0.08.

In the ANC mucus, the levels of MCP-1, Eotaxin (or CCL11), and fibroblast growth factor-basic (FGF-basic) are significantly higher in the elderly than in the young (*p* = 0.033, 0.044, and 0.026, respectively, Fig. 4 and Supplemental Table 4).

**Figure 4.**
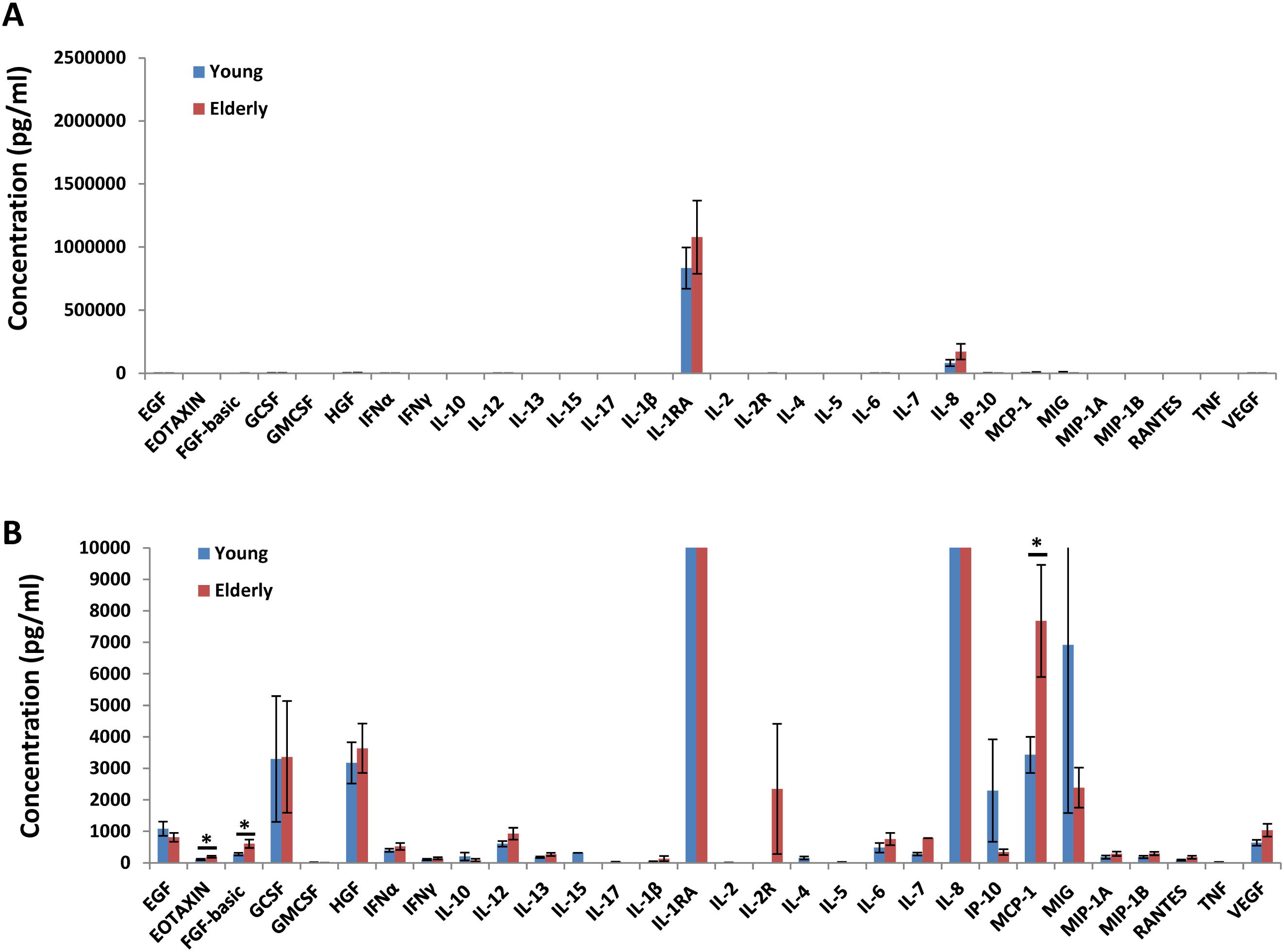
Comparison of cytokine profiles in the ANC mucus from young and elderly subjects. Data were from Luminex multiplex cytokine assays, showing the levels of 30 cytokines and growth factors. (A) Cytokine and growth factor levels showing under large concentration scales to highlight the levels of the two most abundant cytokines, IL-1RA and IL-8. (B) Cytokine and growth factor levels showing in the low concentration range. Data are mean ± SEM. N = 12 per group. * *p* < 0.05.

MCP-1 is a key chemokine that regulates migration and infiltration of monocytes and macrophages (44). MCP-1 can be produced by a variety of cell types, and the source of its production in healthy olfactory mucosa is unclear. In a rodent olfactory bulbectomy model, resident and infiltrating macrophages in the olfactory epithelium appeared to be the major producers of MCP-1 (21, 45). MCP-1 is a chemotactic factor for not only monocytes/macrophages, based on a recent report, it can also stimulate migration of human olfactory ectomesenchymal stem cells derived from olfactory lamina propria (46). The role of MCP-1 in aging of the olfactory system is unclear. To our knowledge, this is the first report of increased MCP-1 levels in the aging human olfactory mucus. Whether MCP-1 is simply a biomarker for increased inflammation during aging or it contributes significantly to olfactory decline during aging remains to be determined. It should be noted that the levels of MCP-1 were higher in both the OC and the ANC mucus in the elderly (Fig. 3 and 4). It has been reported that the circulating levels of MCP-1 increase with aging (47-49). In mice, the gut microbiome composition is associated with increased levels of serum MCP-1 and aging (50). It is plausible that the nasal microbiome composite may also be associated with the increased levels of MCP-1 during aging in both the respiratory and the olfactory mucosa.

IL-12 plays important roles in immunity by regulating T cells and natural killer cells. IL-12 treatments could significantly improve survival in a mouse model of nasal vesicular stomatitis virus infection, which primarily infects the olfactory receptor neurons and then spread through the olfactory nerve to the olfactory bulb and then to the other regions of the brain (51). IL-12 is a heterodimer (p70) composed of a p40 and a p35 subunit. The multiplex assays used in this study detected both the heterodimeric form of IL-12 (p70) and the p40 subunit. Increased levels of IL-12 were detected in the serum of elderly subjects (52). In an Alzheimer’s disease study, high levels of p40 and IL-10 in the blood are significantly associated with amyloid deposition in the brain (53). It is also reported that in mouse Alzheimer’s disease models the levels of p40 in the brain increase with age and inhibition of p40 signaling can ameliorate disease pathology (54, 55). Olfactory decline is one of the earliest symptoms of Alzheimer’s disease (5). Increased levels of IL-12 in the olfactory mucosa during aging might contribute to olfactory loss associated with the neurodegenerative disease.

### The levels of MCP-1, IL-8, IL-13, and VEGF in the OC mucus are significantly and inversely correlated with olfactory threshold performance

To study aging associated olfactory decline, we performed olfactory threshold and odor identification tests. As shown in Fig. 5, threshold sensitivity to PEA and n-butanol was lower among the elderly than the younger adults (Fig. 5A), although the individual tests did not reach significance as analyzed by paired t-test (PEA, *p* = 0.077, n-butanol, *p* = 0.055). However, the combined threshold scores (PEA + n-butanol) did reach significance (*p* = 0.003). Similarly, the percentage of correct odor identification responses were also lower among the elderly, although not significantly (*p* = 0.22, Fig. 5B and Supplemental Table 5)

**Figure 5.**
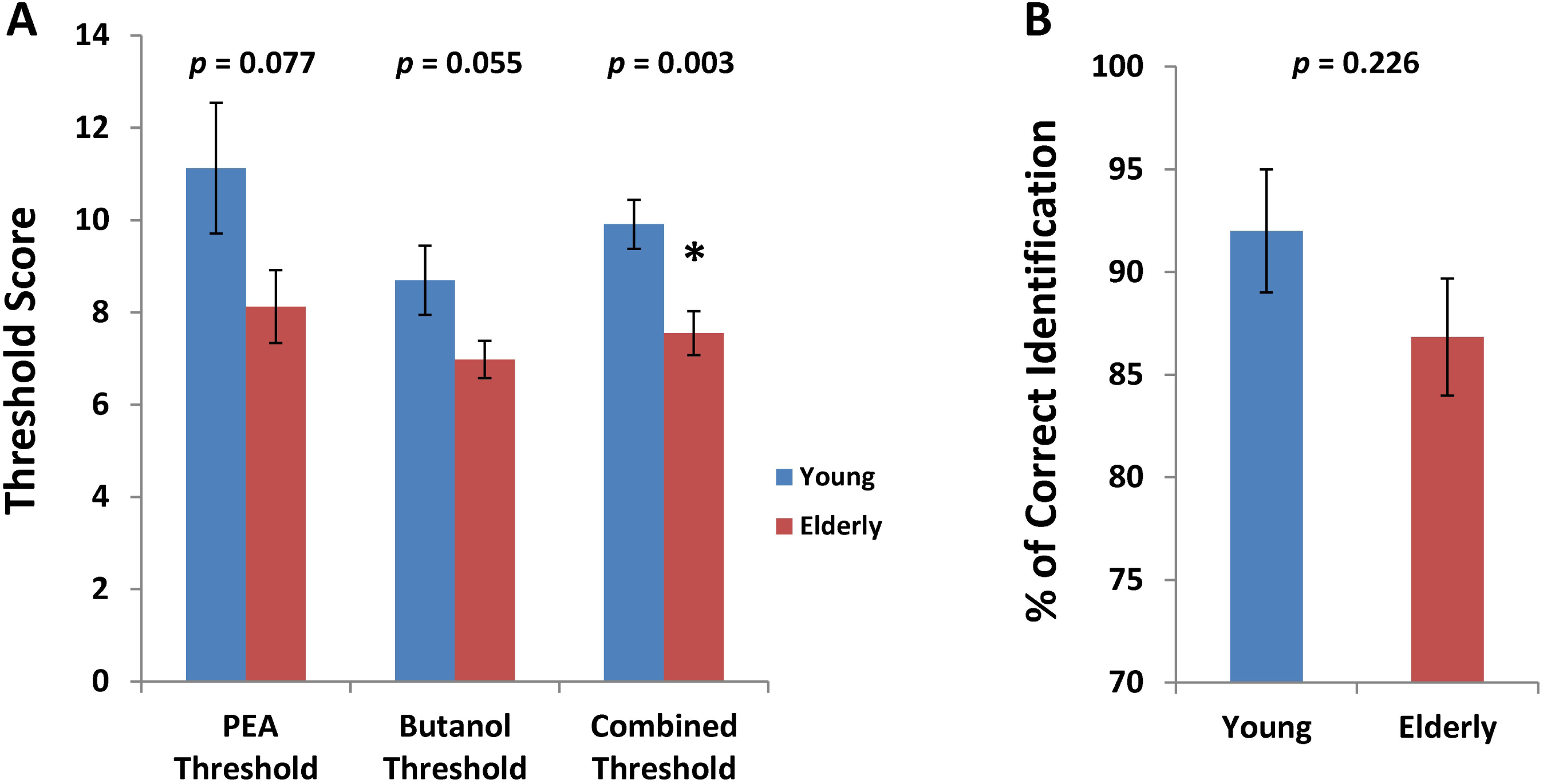
Olfactory performance by young and elderly subjects. (A) Results from olfactory threshold tests, showing threshold scores for phenyl ethyl alcohol (PEA), n-butanol, and the combined threshold for PEA and n-butanol. N = 12 per group. (B) Odor identification tests, showing percentage of correct identification of odor. N = 11-12 per group. Data are mean ± SEM. * *p* < 0.05.

Correlation analyses showed that the levels of four cytokines in the OC mucus were significantly correlated with the combined thresholds of PEA and n-butanol. The levels of MCP-1, IL-8, IL-13, and VEGF were all inversely correlated with the combined threshold scores (the higher the score, the better the performance) (Fig. 6). These results indicate that higher levels of these cytokines are associated with lower olfactory sensitivity.

**Figure. 6.**
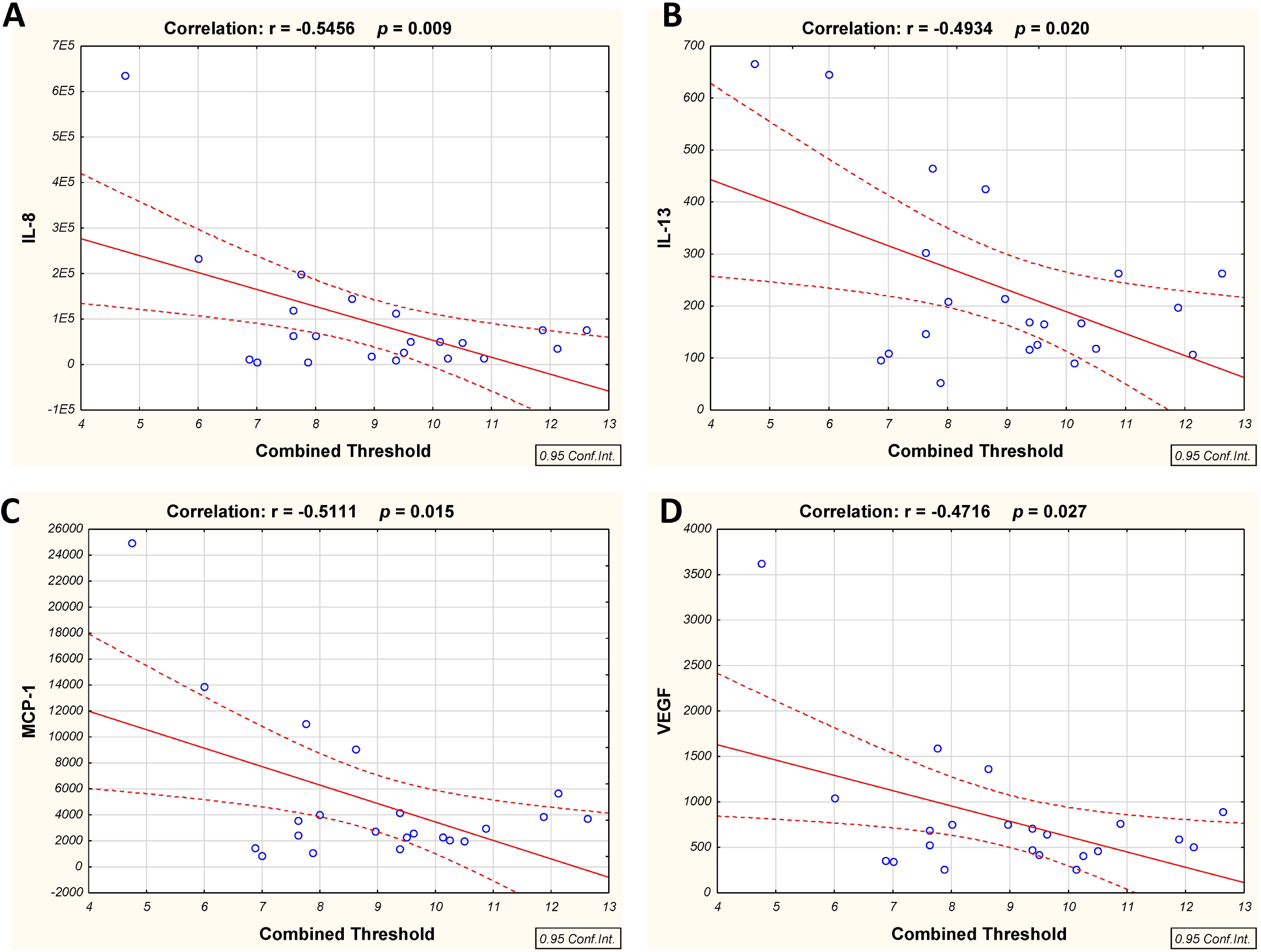
Inverse correlation between cytokine levels and combined olfactory threshold. The levels of IL-8, IL-13, MCP-1, and VEGF, in the OC mucus were significantly and inversely correlated with the combined olfactory threshold of PEA and n-butanol. Y axes are cytokine concentrations in pg/ml. X axes are combined threshold scores. Correlation coefficients (r) and *p* values are as indicated in each panel. N = 22.

All four cytokines are associated with inflammation. MCP-1 and IL-8 are key chemotactic factors for recruiting monocytes/macrophages and neutrophils to the site of inflammation. IL-13 is a T helper 2 (Th2) cytokine that is induced in Th2-mediated diseases, such as allergy (56). VEGF is a potent proangiogenic factor that promotes blood vessel growth during wound healing. Inflammatory responses increase the level of VEGF to stimulate angiogenesis and wound healing (57). The correlations between high levels of these cytokines and low olfactory performance once again indicate that increased inflammation contributes to olfactory decline, consistent with previously published studies. However, whether these cytokines directly contributes to olfactory impairment or they are just biomarkers for inflammation remains to be determined.

## Conclusion

We conducted a study evaluating the factors in olfactory and nasal mucus that may contribute to the observed age-related declines in olfactory acuity. Although we had a small sample of healthy adults (both elderly and young), we observed nearly significant differences in threshold sensitivity for PEA and n-butanol and a significantly lower sensitivity among the elderly when the scores were combined. The cytokine analysis of mucus revealed significantly higher concentrations of specific inflammatory proteins, MCP-1 and IL-12, in the olfactory mucus of older adults relative to younger adults. Of greatest interest, some of the markers observed in olfactory mucus were significantly and inversely correlated with olfactory function. In other words, increased evidence of inflammation in olfactory mucus was associated with decreased olfactory sensitivity. While inflammatory changes have been implicated as contributors to many changes in elderly sensory ability, this is one of the first demonstrations showing a clear correlation between the presence of inflammation in mucus that is secreted from cells in the olfactory cleft and olfactory function.

## Methods

### Human subjects and sample collection

Thirty subjects were recruited by advertisement, and informed consent was obtained from all participants. All procedures were approved by the University of Pennsylvania Institutional Review Board. Mucus samples were collected from 24 adult male and female subjects: 12 younger subjects between the ages of 21-40 and 12 older subjects between the ages of 65-80. Subjects did not have any subjective or objective evidence of sinonasal inflammation based on subject history and nasal endoscopy. Pregnant women were excluded from the study.

Mucus was collected using the sponge part of an eye spear (Medtronic ENT, Jacksonville, FL) that was soaked in double-distilled water and then dried prior to use. Following instillation of topical lidocaine and a decongestant (oxymetazoline) sponges were placed bilaterally under direct visualization: one in the olfactory groove (between the middle turbinate and superior nasal septum for olfactory mucus collection) and one in the other nostril just deep to the nasal vestibule (between the inferior turbinate and inferior nasal septum for anterior nasal mucus collection). After 10 minutes, the sponges were removed. The nasal secretions (mucus) were then centrifuged and divided into aliquots and frozen (58). Mucus collections were done at the Department of Otorhinolaryngology, Head and Neck Surgery of the Hospital of the University of Pennsylvania, and samples were transferred to the laboratories at Monell Chemical Senses Center for processing, storage, and analysis. Aliquots of the mucus were analyzed for cytokine profile by multiplex assays (this report) and global protein contents by shot-gun proteomics approach (see companion article, Yoshikawa et al.).

### Cytokine profiling

A set of 30 factors (Supplemental Table 1), including growth factors, chemokines, and inflammatory cytokines, were analyzed by the highly sensitive antibody-based multiplex cytokine assays (28). The Human Cytokine 30-Plex Panel kit (LHC6003) was purchased from Life Technologies. Mucus samples stored at −80 °C were thawed on ice and diluted with Assay Diluent (from the kit) as suggested by the manufacturer. Due to low volume, two of the olfactory mucus samples from elderly subjects were not included in the analysis. Preparation of cytokine standards and procedure of the assays were carried out following the manufacturer suggested protocol. All incubation steps were done on an orbital shaker with vigorous shaking. Washing steps were done using a vacuum manifold. After the final wash, 100 μl of Working Wash Solution were added to each well. Before reading on a Luminex platform compatible instrument, the beads were resuspended by shaking the plate vigorously on an orbital shaker (500-600 rpm) for at least 2-5 min. The beads were analyzed using a Bio-Plex 200 system instrument (Bio-Rad Laboratories) following the manufacturer suggested settings. The Bio-Plex Manager Software (Bio-Rad Laboratories) was used for instrument control, data acquisition, and data analysis. The software uses Brendan Scientific five-parameter logistic curve fitting model for standard curves. The acquired data were exported into Excel for further analysis and for figure and table preparations.

### Sensory analyses

In order to associate changes in the composition of olfactory mucus with functional outcomes, we conducted sensory tests to characterize olfactory function in our participants. We measured olfactory thresholds for two compounds (PEA and n-butanol) and odor identification for 16 odorants. Sensory thresholds were collected using Sniffin’ Sticks in a 3-alternative forced-choice procedure, as previously described (31, 59). Odor identification ability was also measured using the NIH Odor Identification Test (59), in which individuals are required to choose the correct label out of four possible alternatives. Participants were scheduled for two visits to the Monell Center for the sensory testing. On one visit, thresholds for n-butanol and odor identification were measured. On the other visit, thresholds for PEA were measured.

### Statistical analyses

Statistical analyses were conducted using Statistica version 13 (Dell Inc.) and Excel (Microsoft) software. Data from multiplex cytokine assays were acquired using Bio-Plex Manager software. When the level of a cytokine in a particular sample was below or above detection limits based on instrumental read out, the cytokine in the sample was excluded from analysis. Differences in cytokine levels between groups were analyzed by ANOVA followed by *post hoc t* tests. Differences in olfactory performance between the young and the elderly groups were analyzed by *t* tests. Association analyses were conducted using linear models to identify significant associations between olfactory performance and the levels of cytokines in the olfactory mucus. Combined PEA and n-butanol thresholds were used for these analyses. If the level of a cytokine in the olfactory mucus was below the detection limit in more than 50% of the subjects, the cytokine was excluded from the association analysis. The values from the young and the elderly subjects were analyzed together (N = 22). A *p* value less than 0.05 was considered statistically significant.

### Data availability

All data generated or analyzed during this study are included in this published article and the companion article (Yoshikawa et al.) and the Supplementary Information files.

## Acknowledgements

This research was funded by a Kao Corporation research grant.

## Author Contributions (names must be given as initials)

K.Y., N.S., A.O., J.N., N.C., H.W., and P.D. designed the research project. C.J., N.A., N.C., and P.D. recruited human subjects and collected mucus samples. H.W. and C.J. processed mucus samples and conducted multiplex cytokine assays. C.J. and P.D. performed olfactory tests and analyses. H. W., K.Y., and P.D. analyzed the data and wrote the manuscript. All authors reviewed the manuscript.

## Additional Information (including a Competing Financial Interests Statement)

K.Y., M.H., N.S., J.N. and A.O. are employees of the Kao Corporation (Tochigi, Japan). The other authors had no personal or financial conflicts of interest. Kao Corporation funded the research and made a contract of collaborative study with the Monell Chemical Senses Center.

